# Fast and robust 3D MINFLUX excitation with a variable phase plate

**DOI:** 10.1101/2023.10.31.564945

**Authors:** Takahiro Deguchi, Jonas Ries

## Abstract

MINFLUX has achieved record resolution in superresolution imaging and single fluorophore tracking. It is based on localizing single fluorophores by rapid probing with a patterned beam that features a local intensity minimum. Current implementations, however, are complex and expensive and are limited in speed and robustness.

Here, we show that a combination of an electro-optical modulator with a segmented birefringent element such as a spatial light modulator produces a variable phase plate for which the phase can be scanned on the MHz timescale. Bisected or top-hat phase patterns generate high-contrast compact excitation point-spread functions for MINFLUX localization in the x,y, and z-direction, respectively, which can be scanned around a fluorophore within a microsecond and alternated among different excitation wavelengths.

We discuss how to compensate for non-optimal performance of the components and present a robust 3D and multi-color MINFLUX excitation module with record speed, which we envision as an integral component of a high-performance and cost-effective open-source MINFLUX.

---

MINFLUX^1^ is a super-resolution microscopy technique in which a patterned beam featuring a local intensity minimum, typically a donut beam, probes the signal around a single fluorophore. From the intensities measured at different locations, the position of the fluorophore can be estimated. Recentering of the scan pattern on the estimated position and decreasing its size in an iterative way proves to be a more effective means of enhancing localization precision compared to increasing photon numbers. This means that for a given number of photons emitted by the fluorophore MINFLUX effectively outperforms camera-based Single-Molecule Localization Microscopy (SMLM)^2^ in terms of localization precision. MINFLUX has been used for high-resolution imaging in fixed^3,4^ and live^2^ cells and for tracking single molecules with unprecedented spatio-temporal resolution^5–7^.

For optimal performance, a MINFLUX microscope requires fast and repeated scanning of the pattern to average over potential intensity fluctuations of the fluorophore, fast feedback of the position estimate on the scan pattern, a high contrast of the intensity minimum and a very stable setup combined with active drift stabilization. In most custom^2,5^ and commercial^8^ microscopes, a donut-shaped beam, created by a vortex phase pattern, is scanned laterally using electro-optical deflectors. For 3D MINFLUX, typically a top-hat phase pattern that results in a ‘3D donut’ is employed and axial scanning is performed with an electro-optically actuated varifocal lens^2^ or a deformable mirror^8^. All these fast 3D scanning solutions require expensive components. Standard confocal microscopes with galvo scanners can be upgraded to the MINFLUX mode by insertion of a vortex phase plate^9^, but these implementations might not easily reach the resolution and speed of dedicated MINFLUX instruments. Pulsed interleaved MINFLUX^10^ replaces the lateral scanning with four optical fibers of different lengths at the cost of flexible scan patterns, which in combination with a pulsed laser allow switching between donut positions within 12.5 ns and provides information on fluorescence lifetime. Recently, interference of two beams was used to create bilobed excitation patterns that could be rapidly scanned across a single fluorophore by changing the interference phase with an electro-optical modulator (EOM)^6^. Still, generating a pattern for 3D MINFLUX in multiple colors that features rapid scanning with sub-nanometer accuracy and stability is challenging and can be costly.

Here, we overcome this challenge by developing a MINFLUX excitation module based on a novel variable phase plate, which enables 3D multi-color MINFLUX with high spatio-temporal resolution using a simple, robust, and economic setup.

## Results

### Simple phase patterns for one-dimensional MINFLUX localization

As shown by Wolff et al.^6^, the use of distinctive PSFs for each dimension can outperform the donut-shaped beam profiles generated by a vortex phase pattern. In this work, we use 3 simple binary phase patterns that generate compact MINFLUX beams for x-, y-, and z-localization, respectively (Figure 1 A). For 2D MINFLUX theses bilobed PSFs achieve a similar localization precision compared to donut or interference phase patterns for a given number of detected photons (Figure 1B, Supplementary Figure 1). Due to their small size, they require lower laser powers than the donut beam (Supplementary Figure 2C), which reduces background due to auto-fluorescence and out-of-focus fluorescence, enabling an increase in the density of fluorophores. For 3D MINFLUX the combination of specific patterns achieves a substantially better localization precision compared to using the ‘3D donut’ alone (Supplementary Figure 1).

**Figure 1.**
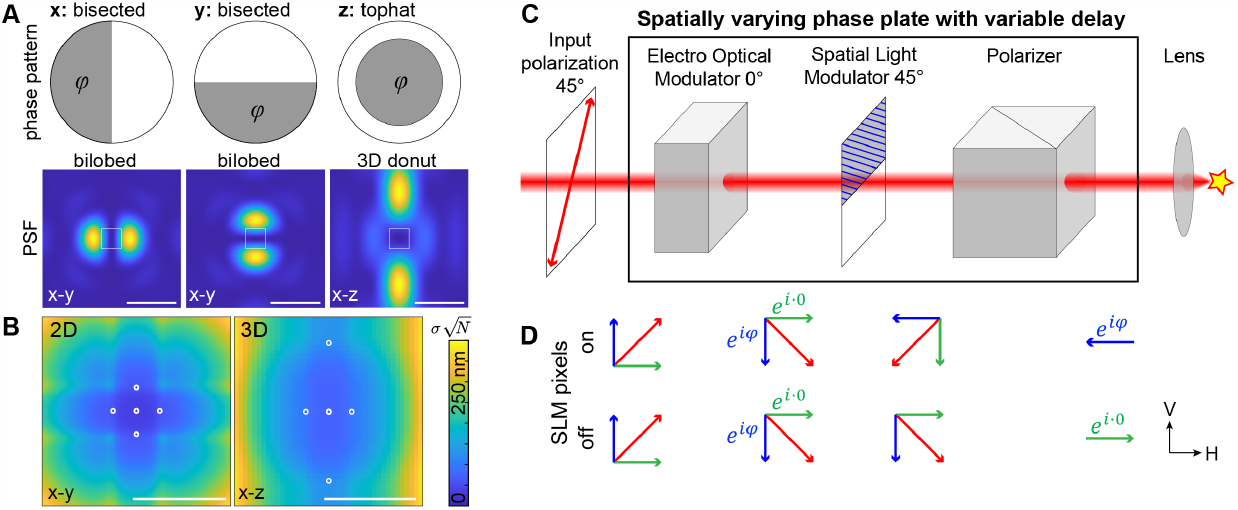
Idea. **A: Phase patterns and point-spread functions (PSFs)** for MINFLUX localization in the x-, y- and z-direction, respectively. The phase *φ* determines the position of the intensity minimum (see Supplementary Figure 3). **B: Theoretical localization precision limit**, the Cramér-Rao bound, for 2D MINFLUX for sequential localization with the bilobed x and y patterns (left) and 3D MINFLUX with the bilobed x and y patterns plus the 3D donut (right), in a region at the centre of the PSF as denoted in A. Simulation parameters: scan range L = 50 nm in the lateral and L_z_ = 150 nm in the axial direction and a background of 0.5% of the maximum intensity when *φ* = 0. The calculated localization precisions are normalized by 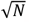, and thus report the localization precision for a single photon. Circles indicate the positions of the minimum during probing. **C: A fast spatially varying phase plate** for MINFLUX excitation comprising an electro-optical modulator (EOM), a spatial light modulator (SLM) and a polarizer. The input beam is polarized under 45° and the output is focused onto the single fluorophore by the microscope objective lens. **D: Illustration of principle**. The vertical polarization component experiences a phase shift of *φ* by the EOM. The ‘on’-pixels of the SLM turn the vertical component into the horizontal component and vice versa, whereas the ‘off’-pixels leave the state unchanged. The horizontal state transmitted by the polarizer thus has the phase *φ* of the EOM for the ‘on’-pixels and no additional phase for the ‘off’-pixels. Scale bars 500 nm (A), 100 nm (B).

Phase differences of π give rise to symmetric patterns with the minimum on the optical axis (x, y pattern) or in the focus of the objective lens (z pattern), because here the π phase shiS leads to destructive interference. Phase differences different from π displace the position of the minimum because these rays must acquire an additional path difference to reach a phase difference of π (Supplementary Figure 3). Simulations show that for the bilobed PSFs a true zero at the minimum is preserved even for large scanning, whereas for the 3D donut the contrast stays high over a z range of a few hundred nm (Supplementary Figure 4).

The phase patterns can be created with a spatial light modulator (SLM), which, in principle, can scan the position of the intensity minimum by changing the phase. However, SLMs that produce continuous phase delays are too slow (<1kHz) for the rapid scanning required for MINFLUX, thus a fast scanner like an electro optical deflector would still be required.

### A fast variable phase plate based on polarization optics

Here we overcome this limitation by combining an EOM, an SLM and a polarizer to generate binary phase patterns with variable phase delay, where the phase delay can be changed on the sub-microsecond time scale by the EOM phase (Figure 1C). The input laser beam is polarized under 45° and enters the EOM for which the polarization axis is oriented vertically under 0°. The EOM can thus cause a phase delay between the vertical and the horizontal polarization component of the beam. The SLM is oriented in a way that the ‘off’-pixels do not change the polarization state, whereas the ‘on’ pixels act as a half-wave plate (HWP) oriented at 45°, which rotate the vertical polarization component into the horizontal polarization. This component carries the extra phase imposed by the EOM. A polarizer (e.g., a polarizing beam splitter) transmits only the horizontal component, which has no extra phase for ‘off’-pixels of the SLM and the phase imposed by the EOM for ‘on’-pixels.

Using Jones matrices, we can calculate in the horizontal/vertical coordinate system that an input beam of 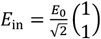 acquires a phase of 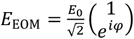 after the EOM. The ‘on’-pixels of the SLM result in 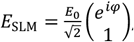, while the ‘off’-pixels do not change the polarization state. After the polarizer the state is 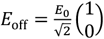 for the ‘off’-pixels and 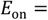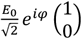 for the ‘on’-pixels. When both components interfere in the vicinity of the focus of the objective lens, the intensity is 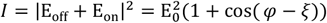. ξ describes an additional phase difference depending on distance to the focal point.

### Non-ideal performance of optical components

The simple setup in Figure 1 is based on ideal performance of the optical elements. Here we discuss the impact of non-ideal performance and experimental challenges and present mitigation strategies.

#### SLM

The binary Ferroelectric Liquid Crystal on Silicon (FLCoS) SLM used here has the birefringent axis of the ‘on’-pixels oriented under 33.5° compared to the ‘off’-pixels, instead of the ideal 45°. Additionally, the phase delay might deviate slightly from π. By calculations and experimentally we found that the addition of a HWP in the beam path before the SLM can perfectly compensate for both imperfections (Figure 2A, Supplementary Figure 5). In addition, the SLM requires ‘balancing’, meaning that for every pixel the time in the ‘on’-state needs to be equal to the time in the ‘off’-state within a 50 ms window. By adjusting the static offset of the EOM phase, the inverse phase map can be matched to the original one, avoiding a reduced duty cycle due to balancing. Finally, the SLM has a finite switching time between different patterns of ∼50 μs, leading to a tradeoff between switching speed and duty cycle.

**Figure 2.**
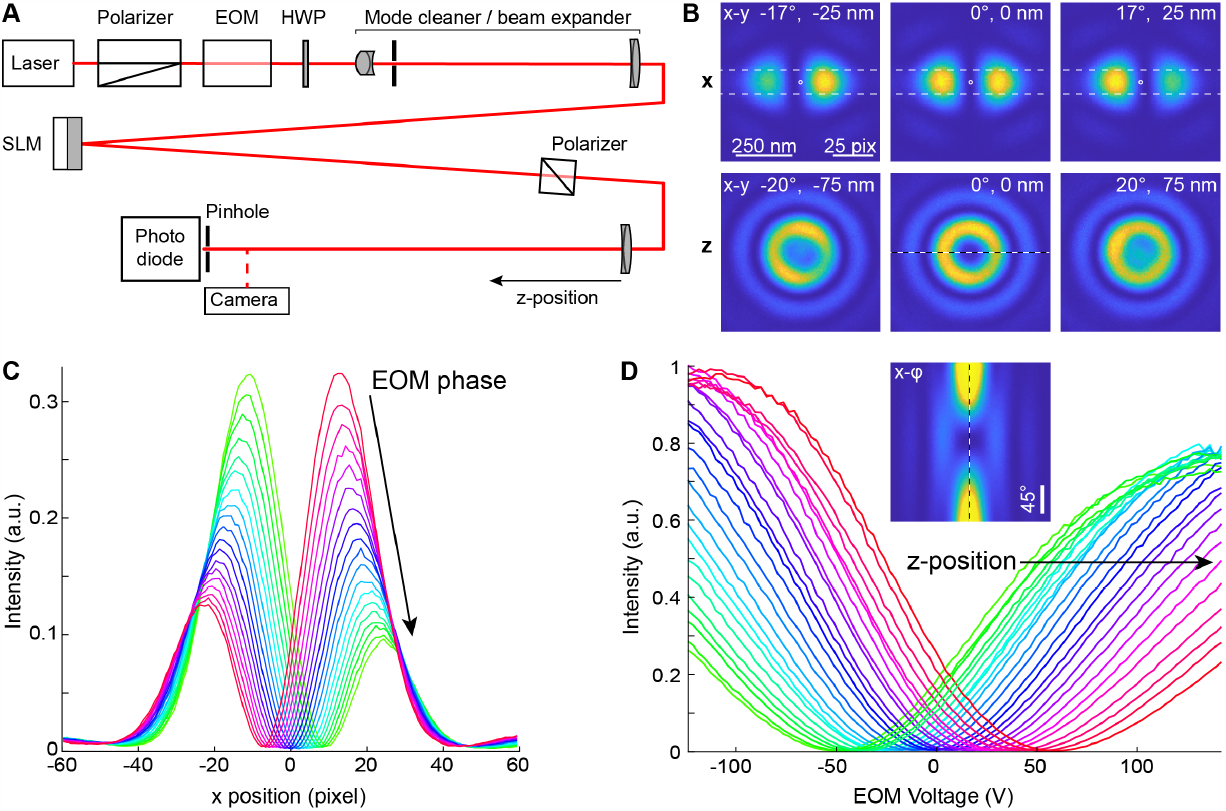
Experimental demonstration with camera detection. A: Beam path of the test setup. The laser is polarized under 45°, passes the electro-optical modulator (EOM), a halfwave plate (HWP) to compensate for imperfections of the Spatial Light Modulator (SLM) and a beam expander comprising achromatic lenses and a pinhole. After the SLM, it passes another polarizer and is focused by an achromat either onto a CMOS camera or onto a small pinhole with a photo diode that detects the intensity a fluorophore would see. **B:** PSFs recorded with the camera for different phases (as indicated) imposed by the EOM for the bilobed x pattern and the 3D-donut z pattern. The scale bar and values in nm are rescaled corresponding to an implementation in a microscope with an NA 1.35 objective lens. **C:** Profiles through the x-PSF as indicated in B for different EOM phases with a difference of 5° between consecutive profiles. **D**. Intensities at the center pixel of the z-PSF for different EOM voltages. Each profile was measured with the last lens positioned at a different distance from the camera (step size 2 mm) and corresponds to measurements with a fluorophore at different z positions (corresponding to a step size of 18 nm in z when integrated in a microscope). Inset: Cross-section through the PSF as indicated in B for different phases *φ* imposed by the EOM.

#### EOM

The EOM phase delay is wavelength dependent, leading to larger displacements of the minima for shorter wavelengths for a given EOM voltage. To compensate this, the static offset of the EOM phase, as well as the amplitude of the phase scan can be set separately for different excitation colors. If necessary, temperature induced phase drifts can be monitored and recalibrated in a separate test beam with a polarizer under 45° and an EOM phase scan around zero phase delay, for instance during dead times caused by switching of the SLM, or with a pinhole and photo diode as in Figure 2A.

#### Silver mirrors

Most mirrors, including silver mirrors, lead to a phase delay between the s and p polarization components, which can spoil carefully engineered polarization states. By aligning the EOM in along the s/p coordinate system and placing the polarizer directly after the SLM, any phase imposed by the mirrors can be compensated by adjusting the static offset of the EOM phase.

#### Aberrated wave fronts

Laser sources, especially free space diode lasers, can have an imperfect beam profile. Additionally, optical components (wave plates, EOM, mirrors) can further deteriorate the wave front, leading to aberrated PSFs and a reduced contrast of the intensity minimum. Here, whenever possible, we place optical components (EOM, waveplate) before the mode cleaner, which then produces a close to ideal wavefront. To reduce astigmatism and coma introduced by a slightly curved SLM, we choose a small beam size on the SLM. In the future, we will insert a second SLM before the microscope to compensate for aberrations from the objective lens or sample.

#### Polarization of MINFLUX PSFs

The x and y phase pattern require the polarization to be parallel to the phase boundary, which can be fulfilled only for one of the directions. One solution is to reflect the beam for a second time on the SLM and rotate the polarization state only for the x pattern (Supplementary Figure 7). The non-ideal direction of the birefringent axis can only rotate the polarization by 67° leading to a calculated deviation of at least 11.5° from the optimal polarization for both the x and y patterns. This results in a contrast of not better than 0.6% (Supplementary Figure 7), which however is still acceptable (see Supplementary Figure 2C). A more accurate alternative is to add a second EOM to the output beam path to generate an optimal polarization state for each phase pattern.

### Experimental demonstration

To experimentally validate our idea without the need of implementing a complete MINFLUX microscope, we designed a beam path in which we replaced the objective lens by an f = 400 mm achromat and the single fluorophore by a small pinhole and photo detector that reports the intensity at the putative single-fluorophore position (Figure 2A). Additionally, we imaged the MINFLUX PSFs with a CMOS camera.

The bisected phase patterns produced the expected bilobed PSFs (Figure 2B,C) with high contrast. By changing the EOM phase, we could scan these patterns laterally with only a minimal loss in contrast (Figure 2C). The top-hat phase pattern produced a high-contrast circular illumination pattern in the focus, and a change in the EOM phase led to an increased signal in the center (Figure 2B). Indeed, when placing the achromat at different distances from the pinhole which effectively changes the pinhole’s z-position, the signal was minimal at different EOM phases, demonstrating that the minimum of the ‘3D donut’ can be scanned in z without degradation (Figure 2D).

To mimic the excitation signal seen by a single fluorophore, we placed a tiny pinhole (0.03 Airy Units) in the focus of the achromat, which transmits the intensity in the central part of the PSF (Figure 3A). We measured the intensity with an amplified photo diode while changing the EOM phase. Specifically, we alternated the x, y and z phase patterns on the SLM and produced a linear EOM phase ramp for each pattern. The intensity trace showed pronounced minima for each phase pattern with an excellent contrast (minimum relative to the intensity detected from a Airy PSF created by a flat phase pattern) of 0.2%, 0.3%, and 0.2 % for the x, y, and z phase patterns, respectively (Figure 3B).

**Figure 3:**
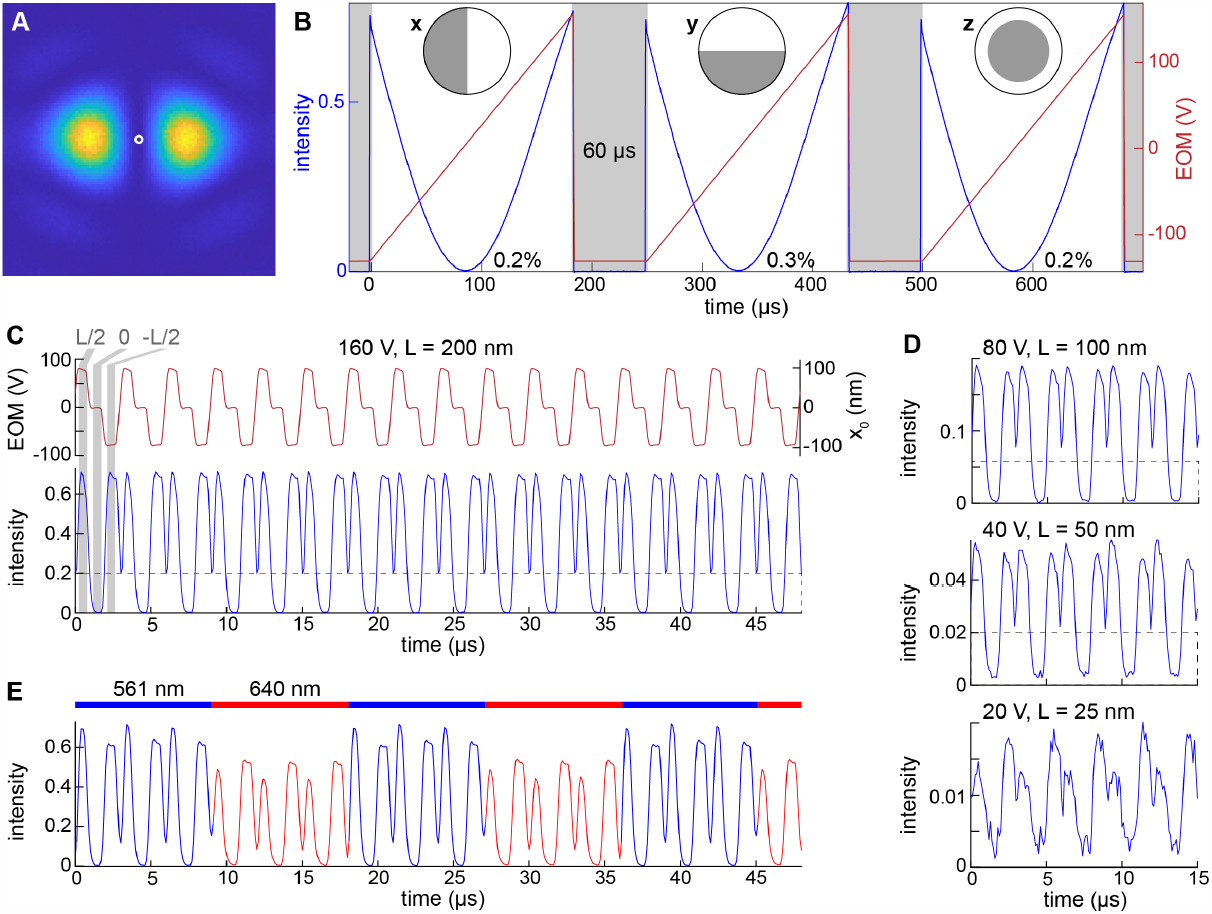
Demonstration of fast scanning using a pinhole, see Figure 2 for the beam path. **A:** An experimental x-PSF acqiured by a camera with the position of the pinhole indicated by a circle. **B:** Intensity (normalized to the intensity measured for a flat phase pattern) recorded through the pinhole for a linear ramp in the EOM voltage. The intensity reaches a minimum (contrast value indicated in the graph) when the minimum of the PSF is at the pinhole position. The measurement is repeated for the different phase patterns (insets) for y- and z-localization, respectively. Switching between phase patterns takes 60 μs. **C:** Mimicking of a MINFLUX measurement. By imposing three different voltages, each lasting for 1 μs, the minimum of the phase pattern is positioned at three distinct positions around the fluorophore. x_0_ indicates the position of the minimum for the case when the setup is used in conjunction with a microsope objective, L is the diameter of the scan pattern. **D:** By changing the amplitude of the voltage, the scan range L can be reduced to improve the localization precision. **E:** By using two colinear laser beams of different colors which are alternated (in this case every 9 μs correspodning to 3 scan patterns), dual-color MINFLUX excitation can be realized without further modifications of the setup.

Next, we tested the speed of our MINFLUX excitation module by repeatedly applying three different EOM phases, mimicking the positioning of the x-PSF at 3 positions around the fluorophore. With excitation times as low as 1 μs per position we could reliably detect three different intensity values with high contrast (Figure 3C, Supplementary Figure 6). Note that our time resolution is currently limited by the electronics for EOM scanning and the bandwidth of the photo diode and can in principle be at least one order of magnitude faster. But even the current implementation is sufficiently fast to not limit any MINFLUX application, as it allows ∼60 scanning cycles within the 180 μs localization time we use for each coordinate.

Multi-color MINFLUX excitation can be realized with the same setup without modifications by using as an input co-linear laser beams. In case of small distances between the fluorophores and low spectral dependence of the phase delay *φ*, both lasers can be used simultaneously. Otherwise, both lasers can be alternated (Figure 3E). As switching between the lasers takes less than one microsecond, the colors can be alternated many times within one MINFLUX localization time, leading to quasi-simultaneous localization of two or more fluorophores.

## Discussion

We developed an excitation module for 3D and multi-color MINFLUX that combines fast and precise positioning of the intensity minimum with a robust and affordable setup. Probing intensities around the single fluorophore takes as little as one microsecond. This enables averaging over fluorophore flickering or a temporally varying background, which otherwise would lead to a bias in the localization. The ability to switch quickly between different phase patterns allowed us to use an optimized PSF for each dimension. The bilobed PSFs that we use for lateral localization result in a high precision for a given number of detected photons (Supplementary Figure 1). They have a smaller footprint compared to donut^1^ or interferometric PSFs^6^, and especially compared to the 3D donut used for 3D MINFLUX^2^, and are thus more robust for higher densities of fluorophores. More importantly, they require lower laser powers, leading to a reduced background and ultimately improved localization precisions (Supplementary Figure 1, Supplementary Figure 2). The high probing speed also allows for more complex scanning patterns with a higher number of positions. It remains to be explored if this can improve the background estimation and localization precision.

Currently, multi-color MINFLUX is performed sequentially with different fluorophores, or using a single excitation laser in combination with fluorophores of slightly different emission wavelengths^2^. This however precludes using MINFLUX for co-tracking of two colors simultaneously. In principle, multi-color MINFLUX could be implemented by duplicating the excitation beam path for each color. With our implementation, however, this is not necessary, as the same excitation module can be used for simultaneous or quasi-simultaneous MINFLUX localization of two fluorophores or more.

As all beams are colinear and are not split up to generate different patterns, colors, or interference phases, they cannot misalign with respect to each other. Thus, our setup is intrinsically robust and stable, which is essential to reach sub-nanometer accuracies in MINFLUX and to use it for routine biological applications. Stability is further supported by the simplicity of the setup with few components and short beam paths. Compared to components employed for pattern generation and 3D scanning in commercial and custom instruments, our solution is very cost-effective, making it a key element for the future development of an affordable open-source MINFLUX instrument with highest performance.

## Methods

### Calculation of polarization states and point spread functions

#### Calculation of polarization states

The polarization state of the beam was calculated in Mathematica (Wolfram) using Jones matrices (**Data and code availability**). Here, a linearly polarized beam is described by

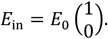

in the horizontal / vertical coordinate system, and a beam with 45° linear polarization by

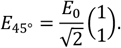

A waveplate (e.g., halfwave plate or SLM) with a phase delay of *φ* and an angle of *α* with respect to the horizontal axis is described by

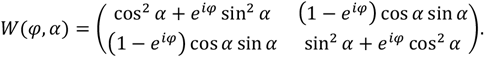

The EOM with a phase delay of *φ* is described by

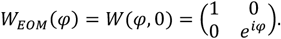

A polarizer transmitting horizontal polarization is described by

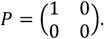

The simple setup (Figure 1) can then be modeled for on-pixels as:

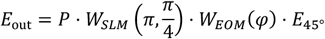

A realistic setup with experimental imperfections is modeled as:

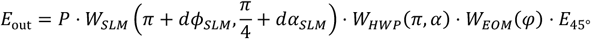

For Supplementary Figure 5 we summed up the field for on- and off-pixels to calculate the intensity after destructive interference. For a given combination of *dϕ*_*SLM*_ and *dα*_*SLM*_ we determined numerically the HWP angle *α* and EOM phase *φ* that minimized this intensity.

#### Calculation of MINFLUX point-spread functions

For the numerical calculations of the electromagnetic field near the focus of an objective lens, we used MATLAB (R2022a, MathWorks) and a software package^11^. We used the following parameters: 100x objective lens, aperture diameter 6.5 mm, numerical aperture 1.35; refractive index 1.406 for the immersion and mounting media; wavelength 635 nm; beam diameter 7.0 mm; flat illumination intensity profile; circular polarization for 3D donut and linear polarization for the rest; and voxel size 2 nm, unless otherwise specified. For the calculation of the interferometric PSF used in Wolff et al.^6^, we used two beams with a Gaussian intensity profile, a beam diameter of 2.3 mm, the beam offset from the center of 2.3 mm, and a relative phase delay of *π* between the two beams^12^.

#### Calculation of Cramer-Rao-Bounds (CRBs)

For calculating the theoretically best possible localization precision for specific PSFs, scanning schemes, detected photons and signal to background ratios we followed Masullo et al.^13^ (Supplementary Figure 1, **Data and code availability**). During MINFLUX localization, the fluorophore at position ***r***_*E*_ is probed with different PSFs *I*_*i*_(***r***_*E*_) to result in *K* emitter intensities ***n*** = [*n*_1_, *n*_2_, …, *n*_*K*_]. This includes the case when the PSF is positioned to coordinates ***r***_*i*_, then *I*_*i*_(***r***_*E*_) = *I*(***r***_*E*_ − ***r***_*i*_), but also the case where the *I*_*i*_(***r***_*E*_) are calculated explicitly. With

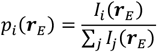

the log-likelihood (after dropping of constant terms) can be written as

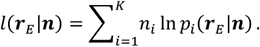

The Fisher information matrix can then be written as:

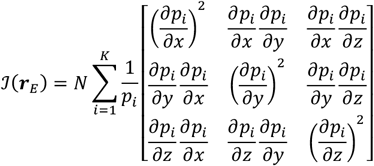

The CRB is then

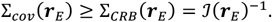

To account for imperfect contrast and a fluorescent background, we modeled the background explicitly by adding an offset to the PSF. To make the background comparable for different PSFs, we calculated the maximum value *I*_A_(0) of an Airy PSF with a flat phase pattern and equal summed intensity and added the offset *b* · *I*_A_(0) to the PSF. We chose this approach over using the signal-to-background (SBR) ratio^13^, because the latter varies strongly across the PSF. The reported maximum localization precisions are normalized to the input photon counts *N* as 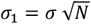 and averaged over the dimensions, with 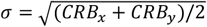 for 2D and 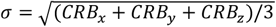 for 3D. They thus report the localization precision of a single photon.

### Optical setup

See Figure 2A. As a light source we used either an iBeam smart 640 nm laser (Toptica) together with an achromatic halfwave plate (AHWP10M-600, Thorlabs) or a multi-color laser engine (MLE HP, Toptica) together with a collimator and beam expander (GBE05-A, Thorlabs) to achieve a beam diameter of approx. 1 mm and a polarization direction of around 45°. The beam was adjusted through a Glan-Thompson polarizer aligned at 45° (GTH10M-A, Thorlabs) and an EOM (EO-AM-NR-C4, Thorlabs) that was mounted under 45° with a custom holder so that its birefringent axis was vertical. As the MgO-doped lithium niobate in this EOM displays a slow negation to an applied DC field for wavelengths shorter than approx. 600 nm, we used a different EOM (LM0202 KD*P 3×3, LINOS) for multi-color measurements, also mounted with its birefringent axis aligned along the vertical direction. The beam then passed an achromatic halfwave plate (AHWP10M-600, Thorlabs) that was used to compensate for imperfections of the SLM and a mode cleaner/beam expander consisting of an f = 40 mm achromat (Thorlabs), a 20 μm pinhole (Thorlabs) and an f = 150 mm achromat (Thorlabs). An iris (SM1D12D, Thorlabs) with a diameter of approx. 2.3 mm resulted in a nearly homogenous beam profile.

The beam was then reflected off the SLM (SXGA-R12-STR, Fourth Dimension Displays) under an angle of approx. 5°, passed a polarizing beam splitter (PBS251, Thorlabs), and was focused onto a 10 μm pinhole (Thorlabs) with an f = 400 mm achromat (Thorlabs). The intensity after the pinhole was monitored with an amplified photo diode (PDA100A2, Thorlabs). Alternatively, the beam was focused onto a CMOS camera (Chameleon CM3-U3-50S5M-CS, Edmund Optics).

### Electronic control

The EOM was driven by a voltage amplifier (HVA200, Thorlabs), which was controlled by a function generator (SDG1062X, Siglent). Intensity signals from the photo diode were acquired with an oscilloscope (SDS2104X-Plus, Siglent). The function generator and the oscilloscope were triggered by the SLM at the start of an image sequence. Additionally, the SLM produces a trigger signal when a valid pattern is established, which was used to switch off the lasers during the pattern switching via a TTL signal. Camera images were recorded asynchronously. For dual-color measurements, we used a custom TTL signal converter in combination with the function generator to alternate between the 561 nm and 638 nm laser line.

### Alignment

We used a polarization analyzer (PAX1000VIS/M, Thorlabs) to align the Glen-Thompson polarizer to 45°. Using a flat phase pattern and large iris diameter, the camera was positioned in the focus of the f = 400 mm achromat. The pinhole was then positioned at an equal distance from the achromat.

Using the top-hat phase pattern, the beam was aligned on the SLM to produce a symmetric PSF. Then the iris diameter and EOM phase were optimized to maximize the contrast of the 3D-donut in focus. The contrast was further maximized by aligning the angle of the half wave plate and the EOM phase.

The pinhole, mounted in an x-y translation stage (ST1XY-D/M, Thorlabs), was adjusted in the lateral directions to maximize the signal on the photo diode.

## Data and code availability

Simulated PSFs, scripts to calculate CRBs and a script to calculate the polarization state are available at: https://github.com/ries-lab/MINFLUXexcitation.

## Acknowledgements

We thank Giuseppe Vicidomini and Eli Slenders for their kind feedback on the manuscript, Luciano Masullo for help with the CRB calculations and the EMBL electronic and mechanic workshops for contributing to the setup. This work was supported by H2020 Marie Skłodowska-Curie Actions (RobMin grant no. 101031734 to T.D.); the European Research Council (grant no. ERC CoG-724489 to J.R.); and the European Molecular Biology Laboratory (T.D. and J.R.).

## Supplementary Figures

**Supplementary Figure 1.**
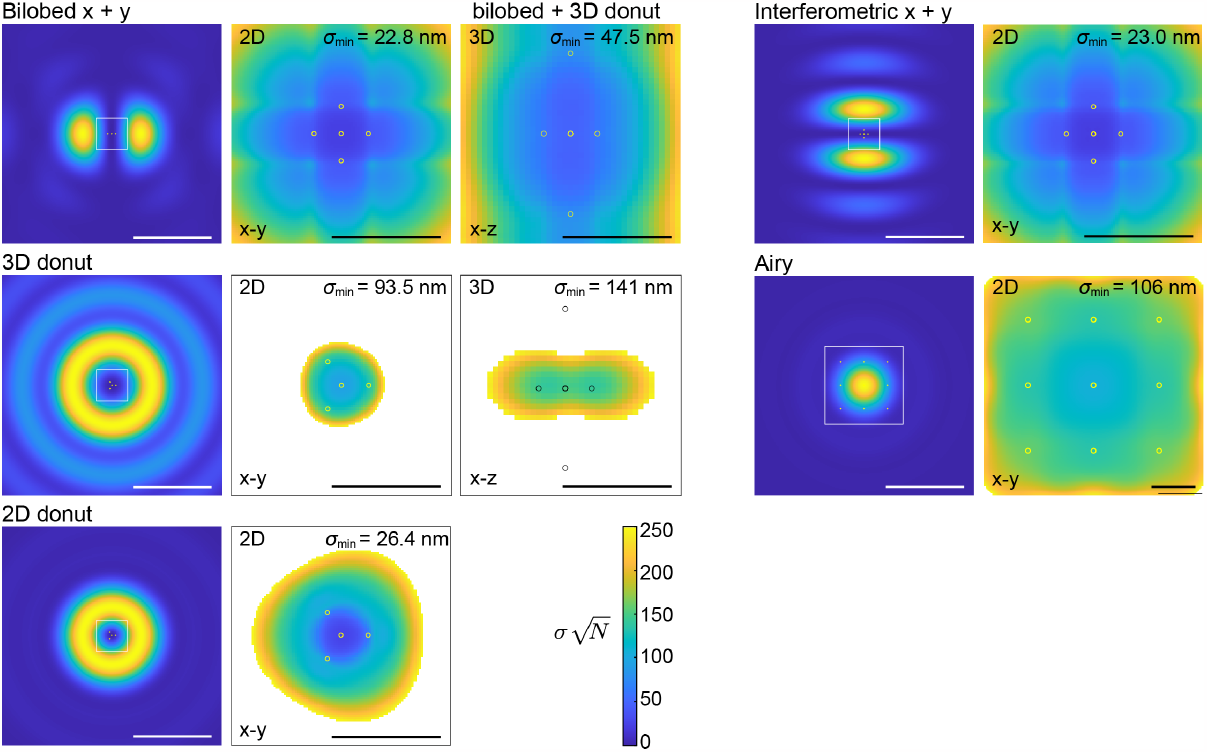
Theoretical localization precision limit for different PSFs. Cramér-Rao bound (CRB) calculations for 2D MINFLUX and 3D MINFLUX with commonly used PSFs. The calculated localization precisions are normalized by 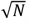, and thus report the localization precision for a single photon. Circles indicate the positions of the minimum during probing. Simulation parameters: scan range L = 50 nm in the lateral and L_z_ = 150 nm in the axial direction, except for the Airy PSF with L = 300 nm for a 3 x 3 scan pattern. Background 0.5% of the maximum intensity of the Airy PSF. Scale bars 500 nm (PSFs), 100 nm (CRB maps).

**Supplementary Figure 2.**
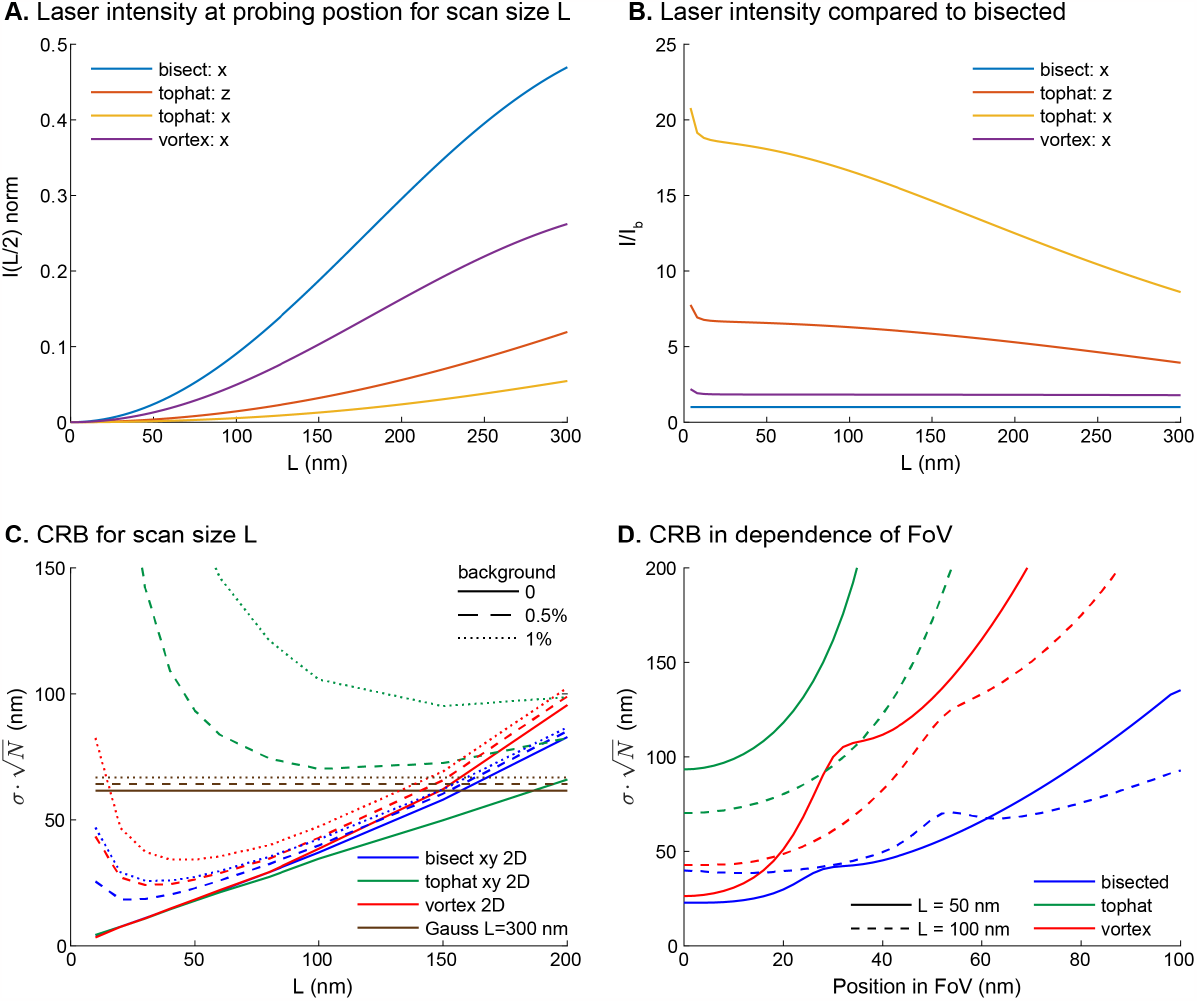
Dependence of localization precision on scan range and field of view (FoV). **A:** Intensity at the probing position for scan sizes L, **B:** Intensity at probing position normalized by intensity for the bisected phase pattern (bilobed PSF), indicating that especially the 3D donut, but also the vortex PSF require higher laser powers for a similar signal. **C:** Maximum localization precision (CRB, normalized by photons) in dependence on scan size L and background for different MINFLUX PSFs, indicating that PSFs with a larger size are more sensitive to background. **D:** Maximum lateral 2D localization precision in dependence on the field of view. Background 0.5% of the maximum intensity of the Airy PSF.

**Supplementary Figure 3.**
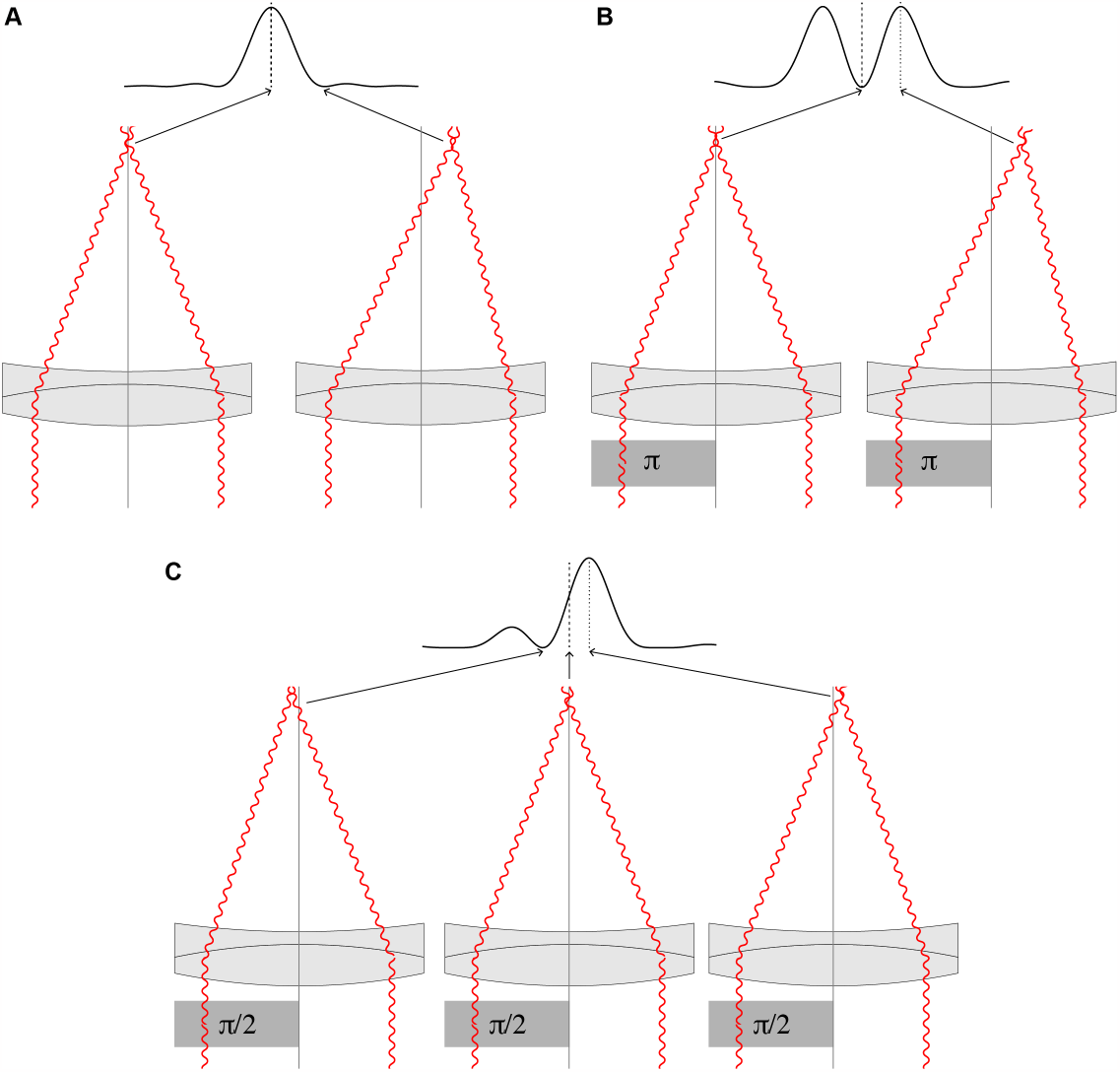
Illustration of how PSFs with a local minimum are generated. **A:** In the focus of a lens on the optical axis, all beams have the same path lengths and thus interfere constructively to result in an intensity maximum. An off-axis point has a longer path length on one side and a shorter one on the other side, leading to destructive interference and an intensity minimum. **B:** A phase shift for half of the beams leads to destructive interference in the focus, and symmetric off-axis with constructive interference, i.e., intensity maxima. **C:** A general phase shift different from π leads to points of constructive and destructive interference at lateral positions that can be chosen by the phase shift.

**Supplementary Figure 4:**
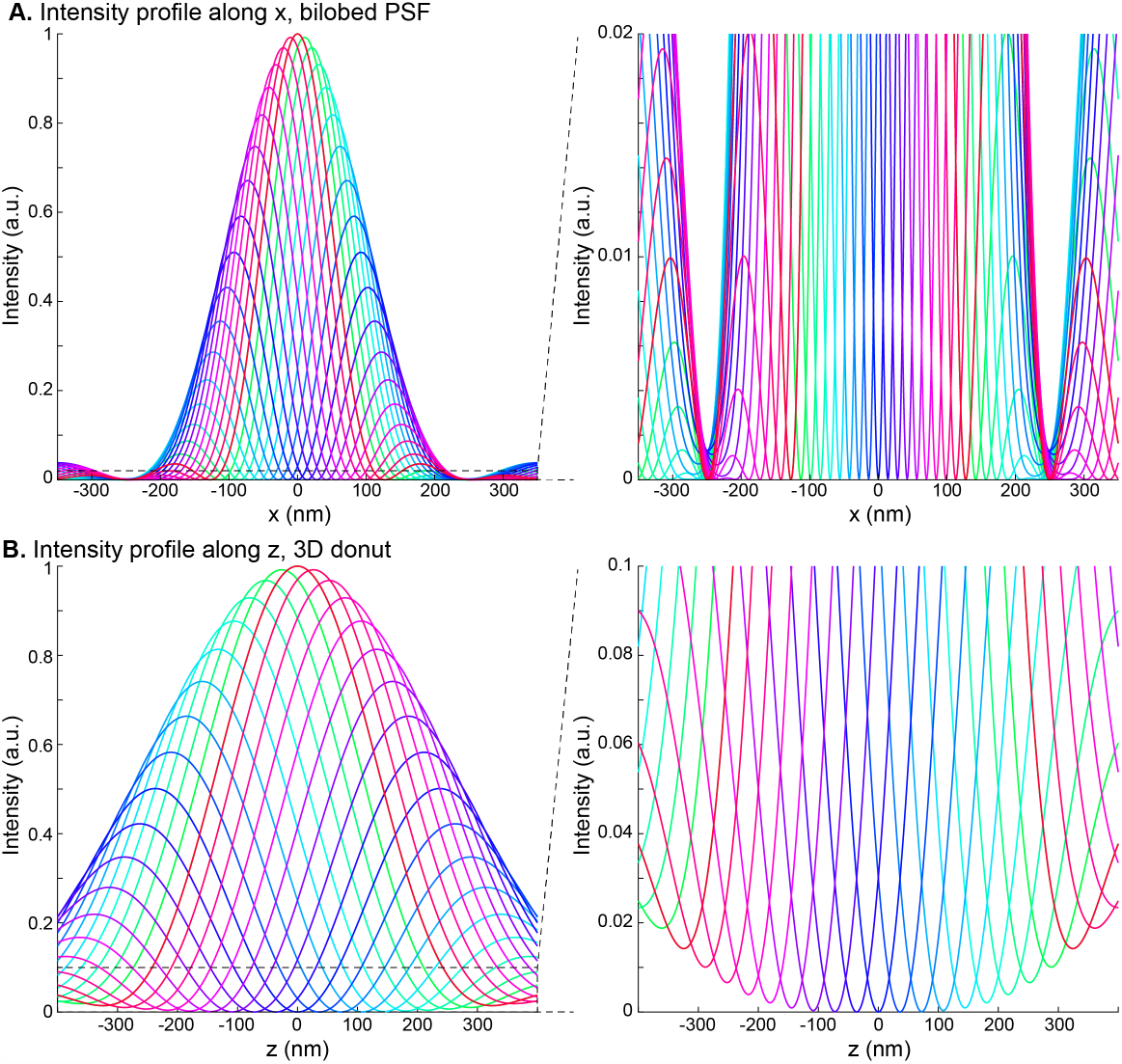
Intensity profiles through simulated MINFLUX PSFs for different EOM phases every 20°. **A: bilobed PSF** from a bisected phase pattern for x localization and zoom into the minima. **B: 3D donut PSF** from a tophat phase pattern for z localization and zoom into the minima.

**Supplementary Figure 5.**
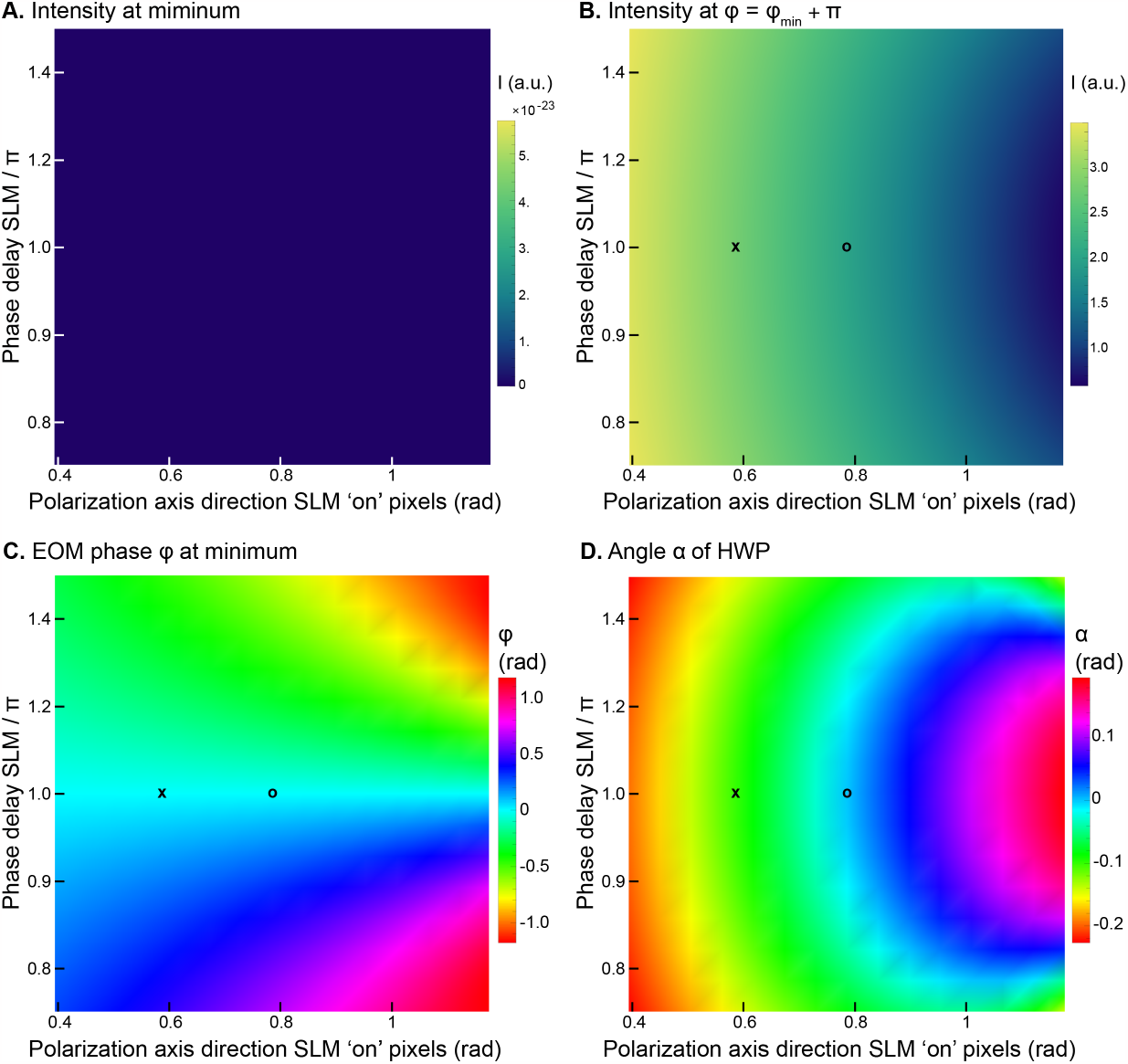
Compensation of SLM imperfections. By inserting an additional halfwaveplate (HWP) in the beam path, we can entirely compensate for an imperfect phase delay ϕ_SLM_ or polarization axis α_SLM_ of the SLM. **A:** We numerically optimized the rotation angle α of the HWP and the EOM phase φ to minimize the intensity in the PSF center and found that for a large range of values the center intensity was zero within the calculation precision. **B:** Intensity for the optimal α for an EOM phase of φ + π as a measure for the efficiency (light transmission) of the setup. **C:** EOM phase φ and **D:** angle of the HWP α for a given combination of SLM phase delay and polarization axis. ‘o’ indicates parameters for an optimal SLM, ‘x’ indicates the parameters in our experiment.

**Supplementary Figure 6:**
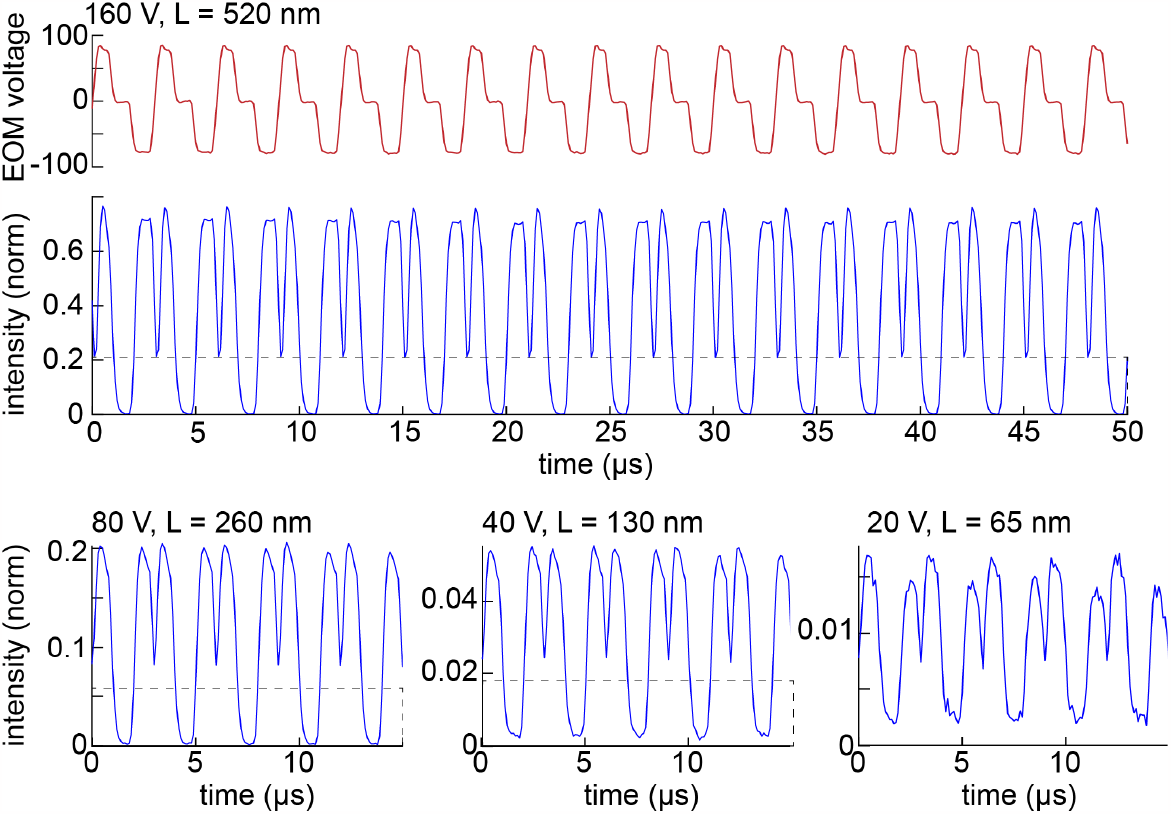
Intensity measurements with a pinhole and photo diode, 3D donut and z scanning. Intensity (normalized to the intensity measured for a flat phase pattern) recorded through the pinhole for three different voltages, each lasting for 1 μs, the minimum of the phase pattern is positioned at three distinct positions around the fluorophore. L is the diameter of the scan pattern when implemented in a microscope. By changing the amplitude of the voltage, the scan range L can be reduced to increase the localization precision.

**Supplementary Figure 7:**
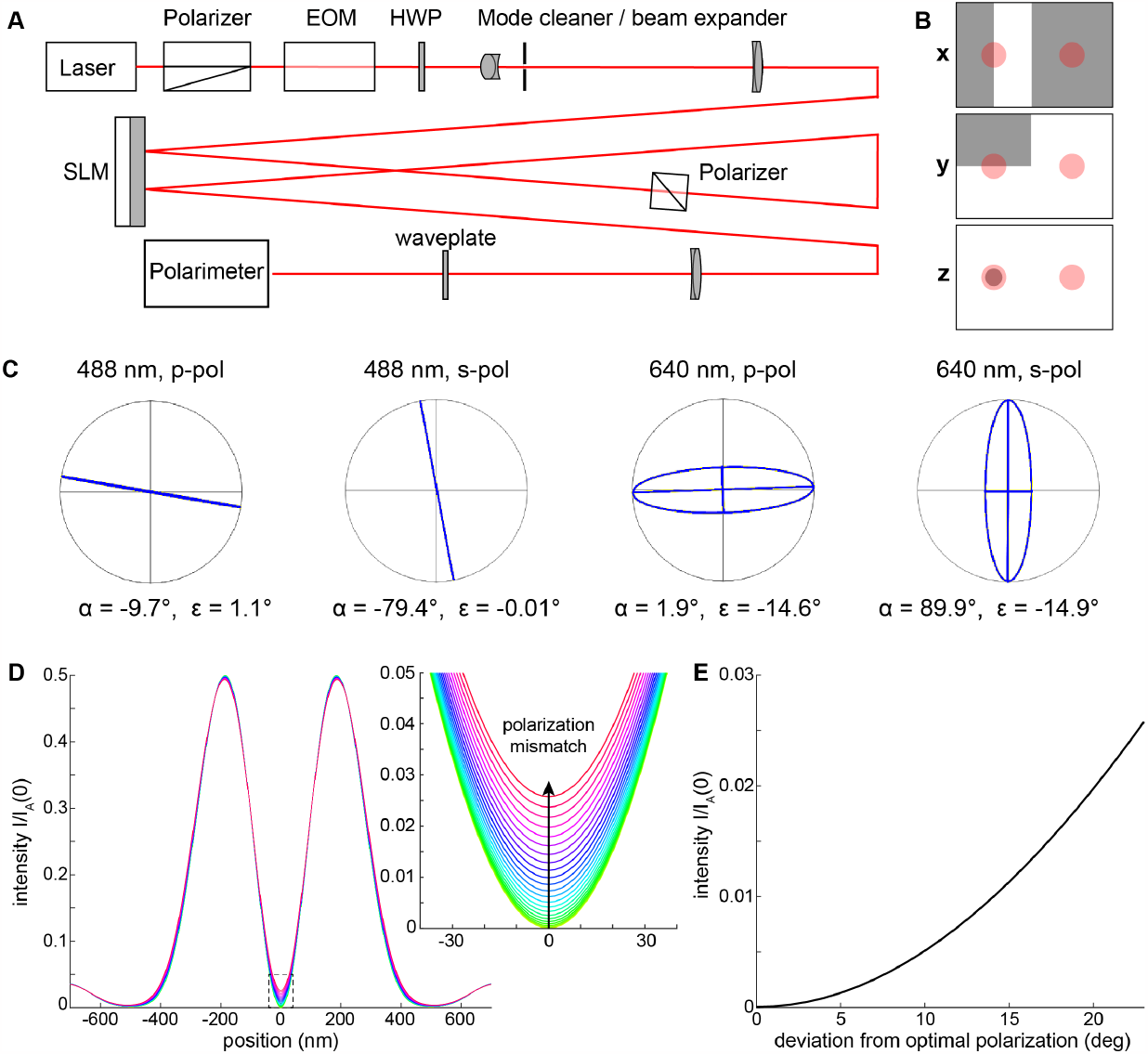
Polarization matching by second reflection off the SLM. **A: Double-bounce beam path** to rotate the polarization of the outgoing beam. **B: Phase patterns on the SLM** for the x, y and z PSFs. The shaded area are the ‘on’-pixels. The red circles indicate the first and second reflection of the laser beam. **C: Measured polarization state** after a waveplate inserted to distribute the polarization error between the two polarization components. **D: Simulated profiles** for the bilobed PSF for an increasing polarization mismatch (1° steps between profiles). **E: Contrast of the PSF minimum in dependence on polarization mismatch**. The intensity is normalized by the maximum intensity of a beam created with a flat phase mask and equal intensity.

